# SISUA: Semi-Supervised Generative Autoencoder for Single Cell Data

**DOI:** 10.1101/631382

**Authors:** Trung Ngo Trong, Roger Kramer, Juha Mehtonen, Gerardo González, Ville Hautamäki, Merja Heinäniemi

## Abstract

Single-cell transcriptomics offers a tool to study the diversity of cell phenotypes through snapshots of the abundance of mRNA in individual cells. Often there is additional information available besides the single cell gene expression counts, such as bulk transcriptome data from the same tissue, or quantification of surface protein levels from the same cells. In this study, we propose models based on the Bayesian generative approach, where protein quantification available as CITE-seq counts from the same cells are used to constrain the learning process, thus forming a semi-supervised model. The generative model is based on the deep variational autoencoder (VAE) neural network architecture.

## 1 Introduction

*Single-cell RNA sequencing* (scRNA-seq) [1, 2, 3] is a powerful tool to analyze cell states based on their gene expression profile with high resolution. RNA sequencing at single-cell level facilitates uncovering heterogeneous gene expression patterns in seemingly homogeneous cell populations. However, the current methods for gene expression profiling at single cell resolution are prone to experimental errors, in particular, inefficient capture of mRNAs [2]. This capture inefficiency results into a general underestimation of the counts (dropout effect). This represents a problem as the current computational approaches for analyzing single-cell data rely on the mRNA counts for clustering and downstream analysis.

Generally, the solution to the dropout problem has been posed as an *imputation* task, where missing counts are filled with estimated counts. Different methods have been proposed for this task, such as non-negative regression [4] or graph-based methods [5]. Another option is to model the dropout effect using the *zero-inflated* (ZI) model [6], where a two-component mixture distribution is constructed, such that the first component models the dropout effect and the second component the observed counts. The effect of *overdispersion* is strongly presented in the scRNA-seq counts, the *negative binomial* (NB) distribution is seen as an appropriate fit to the observed data [7]. Shallow imputation models that are based on zero-inflated negative binomial (ZINB) or zero-inflated log-normal models have been applied to single-cell data [8, 9]. However, these models hypothesize a linear relation between the latent space and the model parameters, which is quite a strong assumption [10]. To overcome the limitations of the linear models, deep neural network architectures have been proposed to resolve missing data (dropouts) [11]. However, discerning technical variation from biological signal solely based on scRNA-seq data is challenging, and assumes that a large number of similar cells are measured.

Accurate imputation strategies are important for downstream analysis, including identification of cell type marker genes, characterization of functional state [12], or the analysis of transcriptome dynamics along differentiation trajectories [13]. An alternative way to approach this problem is to assume that there is a *latent* code that characterizes the cell type (or, more generally, cell state). Conditioning the ZINB distribution with these latent codes would allow sampling accurate transcriptome profiles. In effect, all relevant information about the cell state is included in the latent code. Therefore, the downstream analysis can be based on the model obtained. This approach was proposed by models such as scVI [10] and scVAE [14]. In these techniques and the present paper the goal is to infer the posterior distribution of the latent code [15]. However, the sparseness of scRNA-seq data caused by low mRNA capture efficiency affects the quality of the estimated latents. In order to assess the quality of latent space representations of cell state, manual cell type labeling of the obtained clusters based on marker gene expression has been used. However, this suffers from the dropout effect and introduces a bias related to specific clustering methodology utilized in manual labeling.

Before transcriptome profiling, analysis of surface protein markers has been the mainstream method to decipher cellular identity at single cell resolution. For example, the cellular composition of blood samples has been extensively characterized based on antibodies that recognize specific cell surface marker proteins. Recently, [16] introduced the CITE-seq method that can combine scRNA-seq with such protein marker characterization from the same cells, thus providing complementary data on cell identity. Despite being limited to a small subset of expressed genes, the protein marker count data has the benefit that dropouts are rare compared to mRNA marker gene data. We believed this data could prove useful in assessing the quality of the latent representation. Moreover, it could be incorporated into model training to improve the single-cell model from scRNA-seq [15]. For the SISUA^1^ model presented, we add the protein counts as an additional supervision signal (biological augmentation) with the goal of obtaining higher-quality imputed counts and latent codes. It is noteworthy, that our semi-supervised model uses light supervision in order to assist unsupervised inference instead of the reverse that is usually attempted.

## 2 Single-cell Variational auto-encoding

The task of unsupervised learning is to discover from the observed data [17] hidden structure, such as cluster assignments or low-dimensional representations of the *true data manifold*. In the case of scRNA-seq data, we assume that the true data manifold is of much lower-dimension than the *embedded dimensionality* of the data. Embedded-dimensionality *D*_*e*_ in this case is the number of selected genes in a single scRNA-seq vector x_*i*_ of a cell *i*. A single *batch* of cells has a total of *N* cells and each *x*_*j,i*_ is a non-negative integer, where *j* is the gene index. We denote by *D*_*l*_ the dimensionality of the estimated data manifold. The representation of one cell in the estimated data manifold is typically denoted as a *latent* representation. We will use this terminology in the following text. It is clear, however, that the ultimate goal is to obtain the true data manifold and then conduct inference of the relevant biology from it.

Principal component analysis (PCA) is a classical technique which yields a low-dimensional representation of a high-dimensional dataset. The PCA model can be viewed as a latent representation that minimizes the reconstruction error to generate the embedded space. The reconstruction error is defined as mean squared error (MSE) between the original sample and the generated sample. PCA has been generalized using the Bayesian formalism in probabilistic principal component analysis (PPCA) [18]. In this model, the latent representation is assumed to represent a random variable with prior distribution usually set to 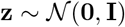. Due to its linear-Gaussian structure, *closed-form* posterior inference is available, and given the latent vector **z** the generation (reconstruction) is also Gaussian

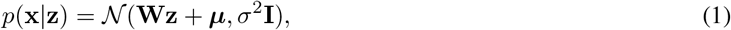

where **W**, ***μ*** and *σ*^2^ are parameters of the model. Estimation of model parameters is performed in the maximum likelihood fashion after latent variables are integrated out, either by closed-form equation or alternatively using the *expectation maximization* (EM) algorithm.

Non-linear models like PPCA are typically not integrable in closed-form, so they require alternative solutions for parameter estimation. In the neural network literature, the autoencoder [19] structure was developed for this task. Autoencoders are deep neural network models that aim to learn the low-dimensional representation, based on a structure consisting of an *encoder* network, which performs the inference, a *bottleneck* layer, which constrains the dimensionality, and a *decoder* network, which performs the generation. The input is fed into the encoder network that converts it into a low-dimensional representation in the bottleneck layer, and the decoder network expands (*reconstructs*) it back into the original signal space. The aim is to reconstruct the input signal with minimal loss, which is typically measured by the *mean squared error* (MSE) function. This learning target correspond to good reconstruction of the input single-cell RNA-seq profile.

### 2.1 Parameterization for the negative binomial distribution

The single-cell gene expression is characterized by raw count data with overdispersion [14, 10, 11]. A well-known practice in the field of statistics has been using the negative binomial (NB) distribution for modeling this kind of events [7]. Furthermore, this strategy has been successfully adapted to the field of deep single-cell mRNA modeling [14, 10, 11].

A conventional approach to parameterizing the NB distribution for an autoencoder uses the mean *μ* ∈ ℝ^+^ and *ϕ* ∈ ℝ^+^ dispersion parameters directly to represent gene expression count data [10, 11].

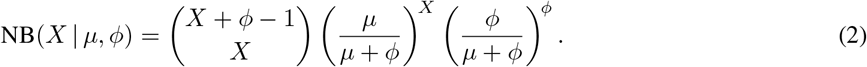

The mean and variance of a random variable *X* ∼ NB(*X* | *μ*, *ϕ*) are

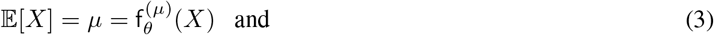

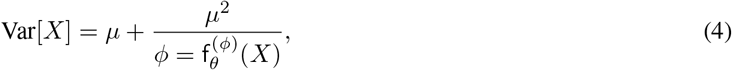

where f_*θ*_ is a deep neural network with parameters *θ*, and the network uses variational techniques and stochastic optimization to estimate *μ* and *ϕ*.

An alternative parameterization is followed by [20], and also implemented in [14]. This approach utilizes two parameters *r* - total count and *ρ* - success rate, which results

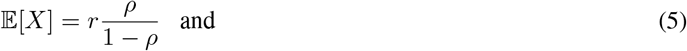

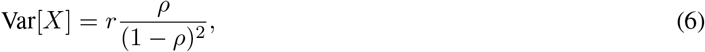

where both *ρ* and *r* also estimated using f_*θ*_. The main difference between Eq. 4 and Eq. 6 is the control of f_*θ*_ over 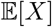 and Var[*X*]:

- Eq. 4 relies solely on the parameter *μ* to estimate the mean 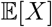. This could turn out to be a high-variance, biased estimate under corrupted data (common to single-cell data sets). In order to compensate for this technical issue, [10] explicitly models cell size, while [11] assumes it is an observed variable provided as an input feature to the network.
- In Eq. 6, f_*θ*_ has more impact since both *ρ* and *r* contribute to the mean and variance of *X*. Since *r* is count values, *ρ* (probability values) could act as learnable scale parameters to adjust the mean of the denoised cell distribution. As a result, this approach implicitly takes into account the cell size during the parameterization.

In this study, we evaluate both approaches, however, we only use Eq. 6 for our semi-supervised extension.

## 3 Biological augmentation using semi-supervised training (SISUA)

The overall design of a multi-output variational autoencoder (*MOVAE*) is illustrated in Fig. 1. Our semi-supervised module is implemented by a linear projection from the decoder output into label space, which is then used to parameterize the distribution of label variable *Y*. The distribution of *Y* could be negative binomial for count data (i.e. NB(*Y* | *r*, *ρ*)) or Bernoulli(*Y* | *ρ*) for probability data. As a result, a comprehensive set of biological information could be directly drawn from the model at once, following this process:

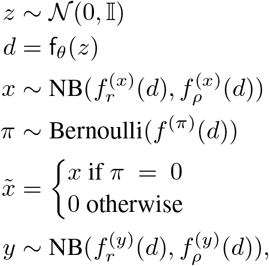

where 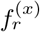, 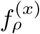 and 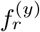, 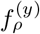 is linear projection function projecting the decoder vector into corresponding dimension for *x* (denoised single-cell gene expression) and *y* (label data such as surface protein expression). *π* is the zero-inflated rate modeled by Bernoulli variables.

**Figure 1:**
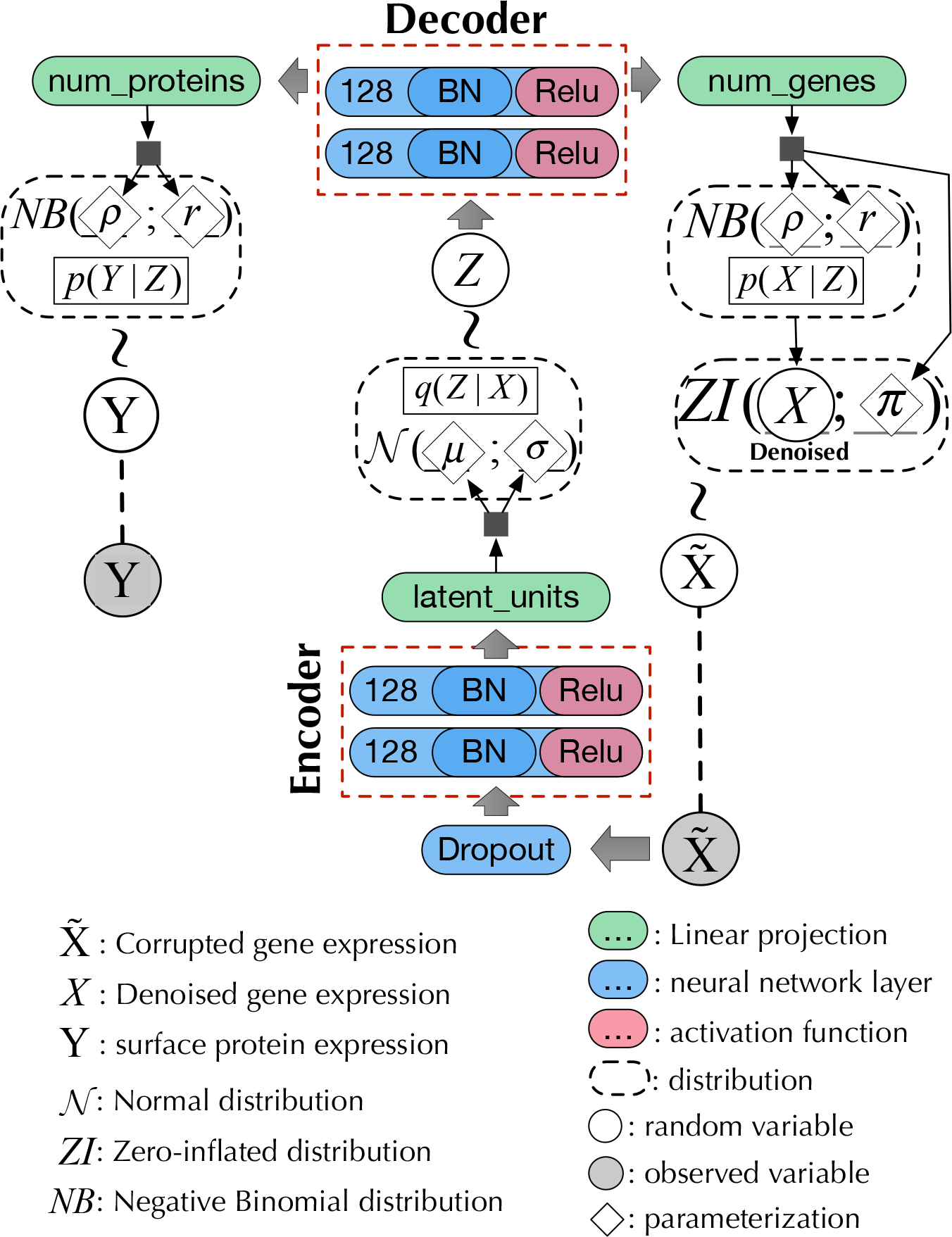
Our variational autoencoder with the semi-supervised extension

### 3.1 Semi-supervised learning for single-cell data

In variational inference, it is possible to have more than one learning target and thereby models that learn a *shared* latent representation. For example, one target could be the reconstruction and another the expressed protein state which could be a probability value, binary value or non-negative discrete measurement. In this manner, learning both tasks together helps to build a more robust representation compared to either task alone. This type of learning is called *multi-task learning* [21]. Multi-task learning can be used in a semi-supervised setting as well, where protein labels are supplied for a subset of the input profiles, which are then modeled jointly to reconstruct mRNA and assign protein marker state, while the rest of the data are modeled only to reconstruct. In the present work, we explore approaches that range from completely unsupervised (no protein labels), semi-supervised (partially labeled) to multi-task (complete label data on protein state) cases.

We further propose three design principles for semi-supervised architectures that improve single-cell gene expression modeling:

- The use of labeled data should only be implicit, i.e. no labeled data should be given as input during the evaluation process. The expense of labeling will typically preclude exhaustively labeled data. As a result, an algorithm, which implicitly encapsulates meaningful patterns from multi-modal data into its latent space during the training phase, would be more robust and practical.
- Enforcing the end-to-end design [22, 23] to avoid the complication of intractable stacked errors, poor scalability to massive data sets, and challenging for practical deployment.
- Unlike conventional semi-supervised learning where an unsupervised objective is created in order to improve the supervised task [15, 24], semi-supervised learning for single-cell data aims for the opposite. Since multiple losses have been known to compete with each other and hinder the major objective of the system [25], Eq. 7 is suggested when incorporating multiple losses into semi-supervised systems.

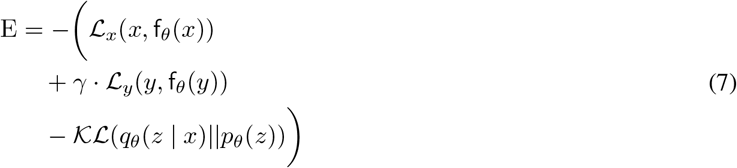

where γ is a hyper-parameter representing the importance of the supervised tasks. Different γ are tested and fine-tuned in Sec. 7. 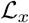 and 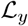 are likelihood functions for the corresponding distributions of the unsupervised and supervised variables. We use ADAM [26], a variation of stochastic gradient descent, to minimize E.

Fig. 2 illustrates the probabilistic graphical model of *SISUA* which satisfies the above design principles, the implementation of which is Fig. 1. The inference process (Fig. 2(a)) parallels the biological relation between mRNA and protein synthesis. The generative process (Fig. 2(b)) enables the sampling of both gene expression and protein marker levels from a biologically-motivated latent space. The implementation of 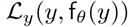 is a major difference between *SISUA* and *MOVAE*. *SISUA* leverages probabilistic embedding to regulate the amount of information backpropagated from the supervised objectives, which will be discussed in the next section.

**Figure 2:**
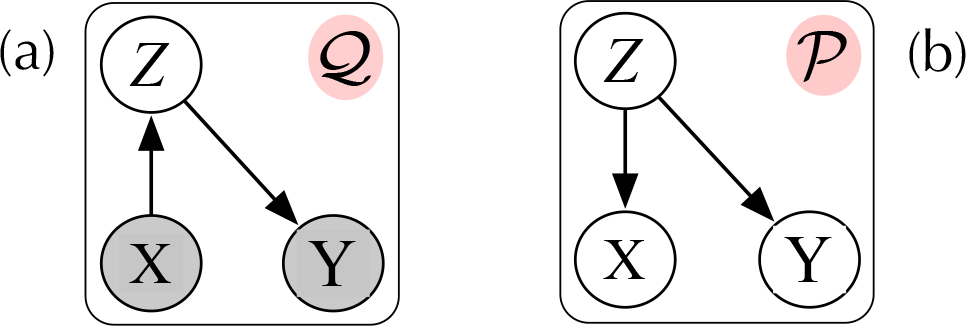
The design of the semi-supervised systems and their probabilistic graphical models. 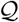 is the inference model and 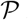 is the generative model.

### 3.2 Probabilistic embedding for biological data

Inspired by recent advances in the field of biometric verification systems [27], we propose a generalized approach for incorporating multi-modal biological data into the unsupervised algorithm. A Universal Background Model (UBM) is a Gaussian Mixture Model (GMM) used to represent general, cell-independent feature characteristics. In our case, the UBM is used to capture different modes of protein activation based on the surface protein levels. Then the model can be used to compare against a cell-specific protein level when making the decision in the presence or absence of a particular protein.

Two considerations motivate the application of the UBM in biological data:

- The data often come from sources (e.g. different measurements) with different characteristic scales and technical variability. These issues pose a different sort of challenge for modeling and possibly hinder the main goal of our algorithm.
- The distribution of the data is often skewed and imbalanced. For example, Fig. 3(c) indicates abnormally high abundance of ‘CD45RA’. This could trigger the false perception that “everything is CD45RA” during the optimization of the deep neural network since the dominant classes will backpropagate most of the updates [28, 29].

For each protein, a two-component GMM is trained,

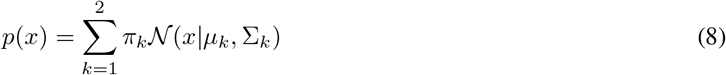

where *x* is a single dimension protein level. *x* could be the raw value in Fig. 3(a) or the log-normalized value in Fig. 3(b). Our experiments have shown that a log-normalized value encapsulate more informative structure of the underlying protein distribution and is less likely to be affected by technical errors and outliers. *μ*_*k*_ and Σ_*k*_ are the mean and covariance vectors of the Gaussians, *π*_*k*_ is the mixture weight. Those parameters are the maximum likelihood estimates [27] that best match the protein distribution.

The UBM utilizes a set of GMM associated with each protein. These GMMs are used to generate the response of the protein to each individual cell, “probabilizing” protein expression. They can be thresholded as in Fig. 3 to yield binary variables indicating the presence or absence of protein in each cell. The final distribution of those process is illustrated in Fig. 3(c), which shows that the binarized and probabilized values are more balanced than the original distribution.

## 4 Experimental setups

We discuss the selection of data sets and the configurations of the baselines and *SISUA* in this section. The experiments were run on two data sets.

The first data set, PBMC, consists of 12039 human peripheral blood mononuclear cells, generated using the 10X Genomics platform. This data set includes cell type labels assigned by manual examination of clusters [30].

The second data set, peripheral blood CITE-seq data, was downloaded from 10x Genomics^2^. Protein marker levels were available for a total of 14 specific antibodies and three control (IgG) antibodies. Here, we utilized the entire dataset or its subset (LY) with markers (CD3, CD4, CD8, CD56, CD16, CD19, CD25, CD45RA, CD45RO, PD1, TIGIT and CD127) that allow distinguishing different lymphoid cell populations (4697 cells, 2000 most variable genes). In addition to raw counts for mRNA, CLR-normalized ADT counts were used for model evaluation and training.

In order to evaluate the generalizability of each algorithm, we split each data set into disjoint training and testing subsets, containing 90% and 10% of the data, respectively. For imputation benchmarking, we measure the robustness of the algorithm by corrupting the original training data, then using the learned algorithm to provide denoised gene expression. Binomial data corruption was applied as in [10]. 25% of the matrix entries are randomly selected and replaced with a Bin(*n,* 0, 2) random variable, where *n* is the original count of the given entry.

Three unsupervised baselines were selected for comparison to *SISUA*:

- Deep count autoencoder (*DCA*) is a denoising autoencoder that takes the count distribution, overdispersion and sparsity of single cell data into account [11].
- Single-cell variational inference (*scVI*) is a framework using deep probabilistic inference to model observed expression values, accounting for the technical variability of the measurements [10].
- *scVAE* [14] also utilizes deep generative modeling. This approach was arguably the least sophisticated because the cell size (or library size) is implicitly modeled via parameterization of NB Sec. 2.1.

**Figure 3:**
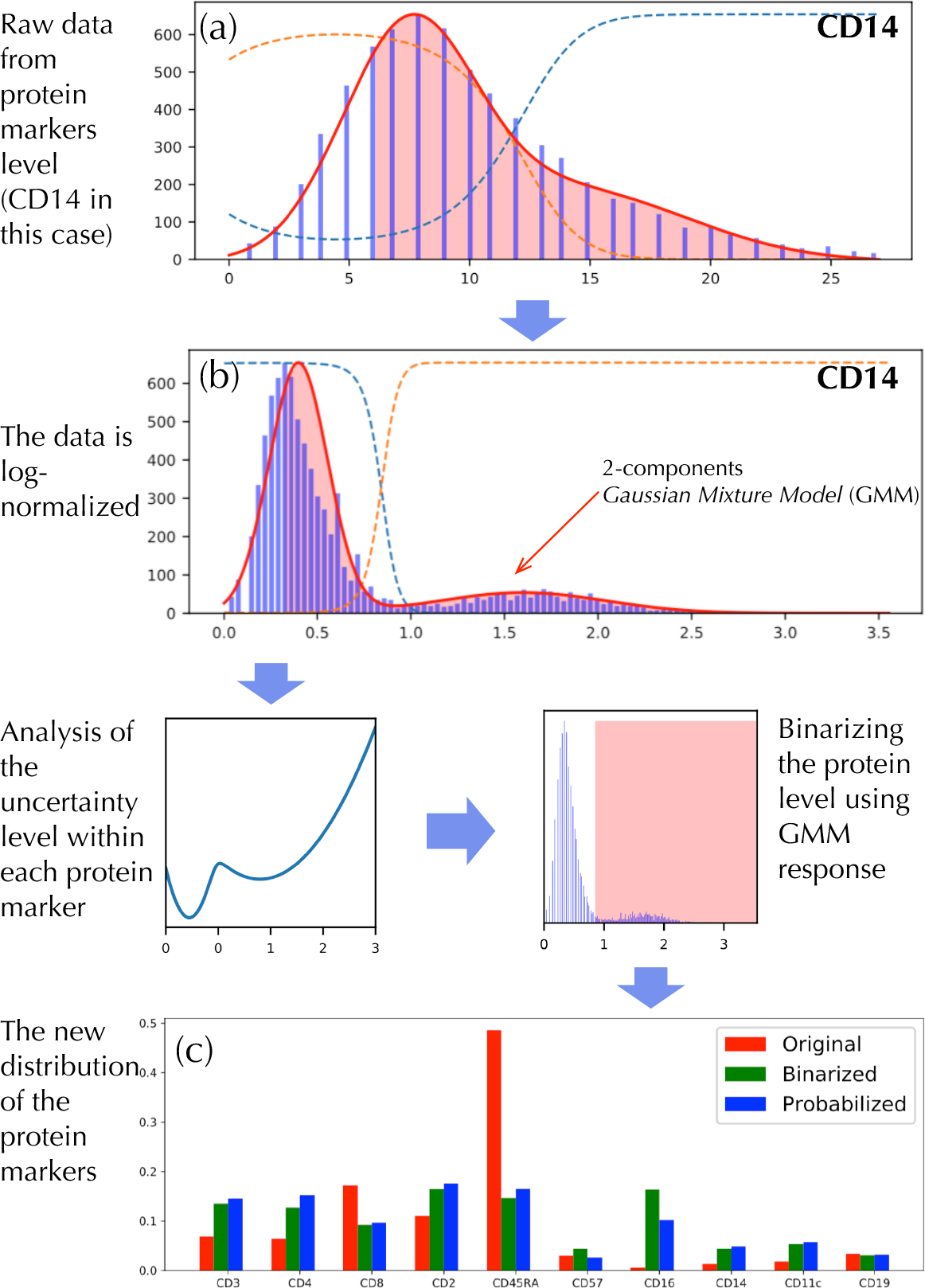
The process of probabilistic protein embedding using data for protein *CD14* as an example. The comparison between the distribution of protein markers before and after the process is shown in the bottom figure.

These three studies represent the state-of-the-art in modeling of single-cell RNA sequencing. We configured each framework similarly to *SISUA* as in Fig. 1. Additionally, the same optimization algorithm and training parameters (number of the epoch, batch size, learning rate) were used in all models.

## 5 Semi-supervised learning enhances biological properties

In this section, we propose experiments to reflect how well the models capture different biological properties of the cell types analyzed. In the following experiments, we focused on three major functions of an auto-encoding model:

- The output space, the denoised gene expression profile, is evaluated using i) per-cell marker protein levels (PBMC CITE-seq), or ii) per-cell assigned labels from the manual examination of data (PBMC RNA-seq) as ground truth for cell types present.
- The latent space, as a low-dimensional representation of the data, is evaluated for biological tasks including inspection, annotation, and recognition of different cell types.
- The semi-supervised space, expressing the protein marker levels (or cell type labels), is a unique feature of *SISUA*. We evaluate the soundness of this space by evaluating its connection to the output and latent space and utilizing the ground truth labels.

**Figure 4:**
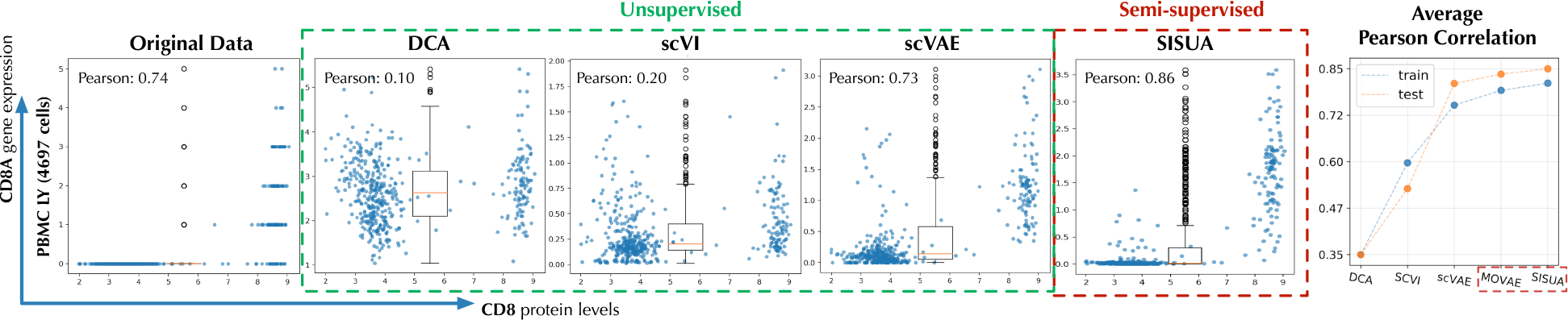
Correlation between CD8A gene mRNA and protein levels in *PBMC Ly* lymphoid cells. The average Pearson correlation for all marker gene/protein pairs is shown in the panel on the right. Semi-supervised models are highlighted by red dashed boxes.

### 5.1 Correlation of marker mRNA gene expression and surface protein levels

Because assaying marker protein levels is less prone (for technical reasons) to the dropout issues that plague mRNA levels for the corresponding genes, cell surface marker protein expression can be used as “ground truth” for evaluating known cell states and cell types. Thus, the denoised corresponding mRNA levels for the same markers can be evaluated in an unbiased manner [16, 11] (Fig. 4).

As exemplified by the T-cell marker CD8, non-zero counts are observed when protein levels are high. This correlation is poorly modeled by *DCA* and *scVI* that impute counts also to cells with low surface protein levels, while scVAE preserves the correlation. Semi-supervised learning consistently improved the correlation across all marker gene and protein levels. In all cases, *SISUA* was able to restore missing gene expression that is biologically plausible.

### 5.2 Separation of cell types in latent space

A trained encoder from each algorithm projects gene expression into low-dimensional latent vectors. To visualize these, the vectors were transformed into 2-D representation by the t-SNE algorithm. A biologically informative latent space should have a clear separation between different cell types. To evaluate this, the points on the t-SNE maps are colored by their ‘ground truth’ labels (manual labels in Fig. 5 and protein marker state in Fig. 6).

Based on pre-assigned labels (Fig. 5) all methods capture a biologically meaningful latent space to a certain extent. Clusters of different cell types are clearly separated. However, many outliers are present in the *DCA*, *scVI* and *scVAE* results and *scVI* shows extra confusion among cells by grouping many clusters close together. It is notable that most of the outliers were classified as ‘Other’ cells. In this label-supervision scenario, *SISUA* yields cleaner cluster structures with fewer outliers, and better minimizes intra-class and maximizes inter-class variance. Moreover, there is a subtle group of ‘NK cells’ placed adjacent to the ‘CD8 T cells’ (SISUA figure panel). This *rightly* calls into question the mutually-exclusive labeling of cells because there, in fact, exist ‘NK cells’ which are also ‘T cells’. Immunologists recognize an entity called NKT-cells as a separate cell type. Among the models, only the latent space representation of *SISUA* clearly indicates the prominent characteristics of the ‘T cells’ within this small group of ‘NK cells’. Notably, this similarity is learned without extra information about the ‘T cells’.

Fig. 6 delivers further insight into the capability of *SISUA* for representing biologically meaningful sub-structure. In *PBMC Ly*, each cell is characterized by multiple protein levels, which include markers for similar cell types (such as CD4 or CD8 positive T cells) that often get tangled up in the latent representation of mRNA data. Compared to all unsupervised representations, *SISUA* achieves strong separation between ‘CD8‘ protein and ‘CD4’ protein in its latent space. This division is biologically plausible and the algorithm is able to learn this pattern independently without explicit cell type labels.

The design of the *SISUA* model only allows an indirect influence of protein markers on the latent space via the supervised objective. It is also possible that this could induce ‘adversarial’ behavior of the learning algorithm [25], hence, defeat the main goal of improving the latent representation by coordinating various biological sources. We did not observe this examining the latent space. Additionally, we evaluate the learned latent spaces quantitatively using two different approaches. As a first method, we feed the learned latent to a secondary classifier that is trained to classify protein markers or cell labels. The results shown in Fig. 7. We notice that in terms of the classification accuracy (F1-score), SISUA is clearly the best model for the training and test portions. The testing performance of the semi-supervised models is degraded when compared to the training portion, but still models clearly win over the fully unsupervised variants.

**Figure 5:**
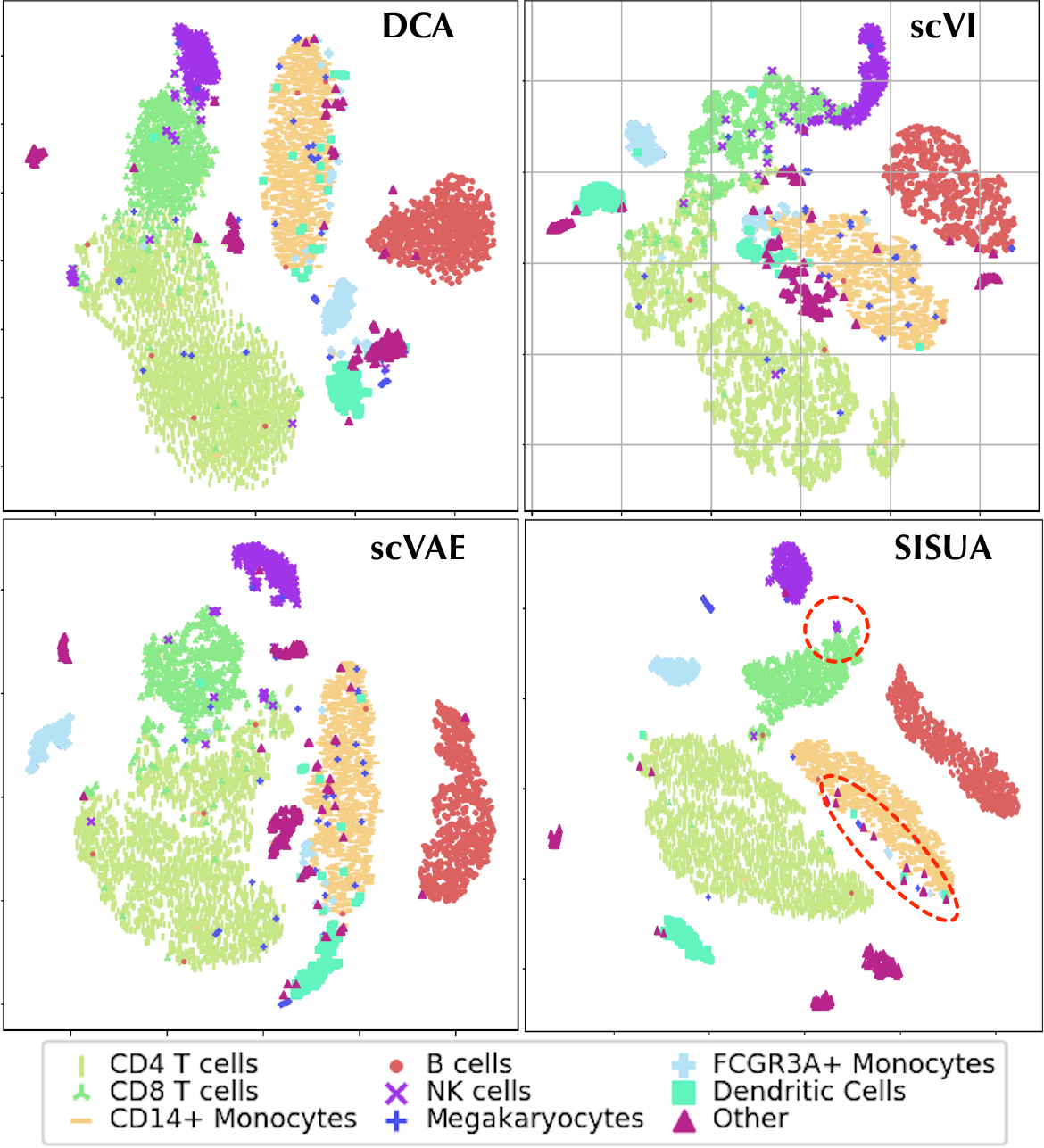
t-SNE visualization of the latent space for *PBMC 10x* dataset, binary cell type labels are used for coloring.

**Figure 6:**
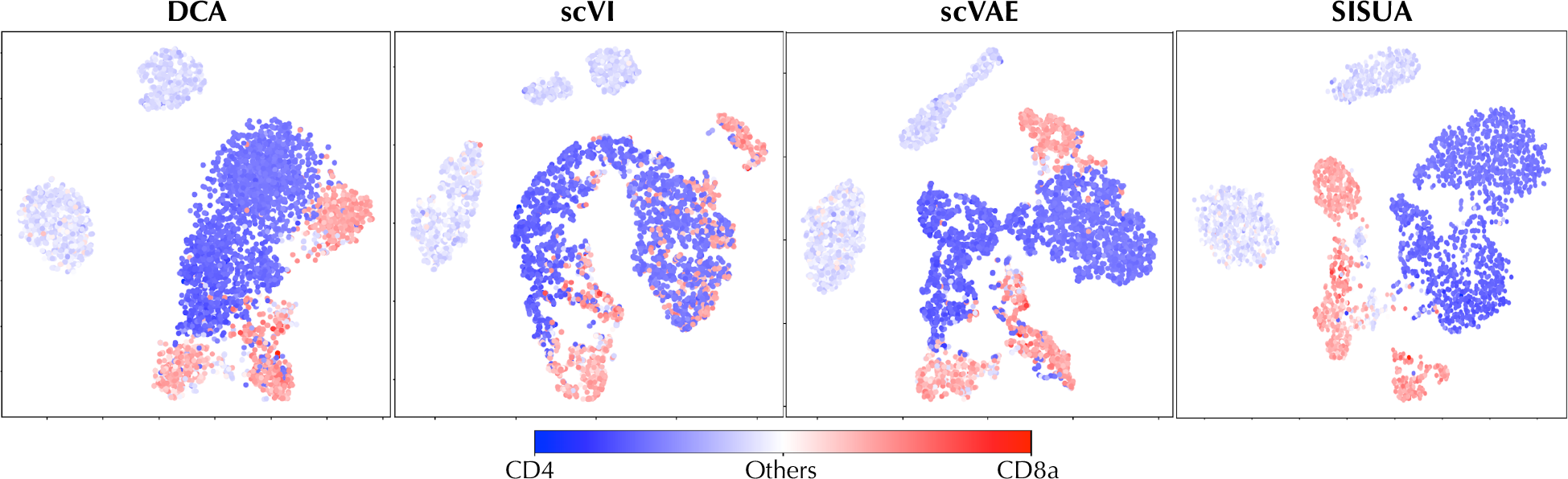
t-SNE visualization of the latent space for PBMC Ly dataset, the activation levels of protein CD8a (dark red tones) and CD4 (dark blue tones) are shown as heatmap.

Noting that we assess usefulness of the learned unsupervised latent code, we turn to classical unsupervised metrics. As a second method, we pool the external validity indices used to asses clustering quality, namely: *ARI* - adjusted rand index, *ASW* - silhouette score, *NMI* - normalized mutual information and *UCA* - unsupervised clustering accuracy.. These results are also shown in Fig. 7. The results indicate that the information encapsulated in semi-supervised latent space is higher than the other models.

**Figure 7:**
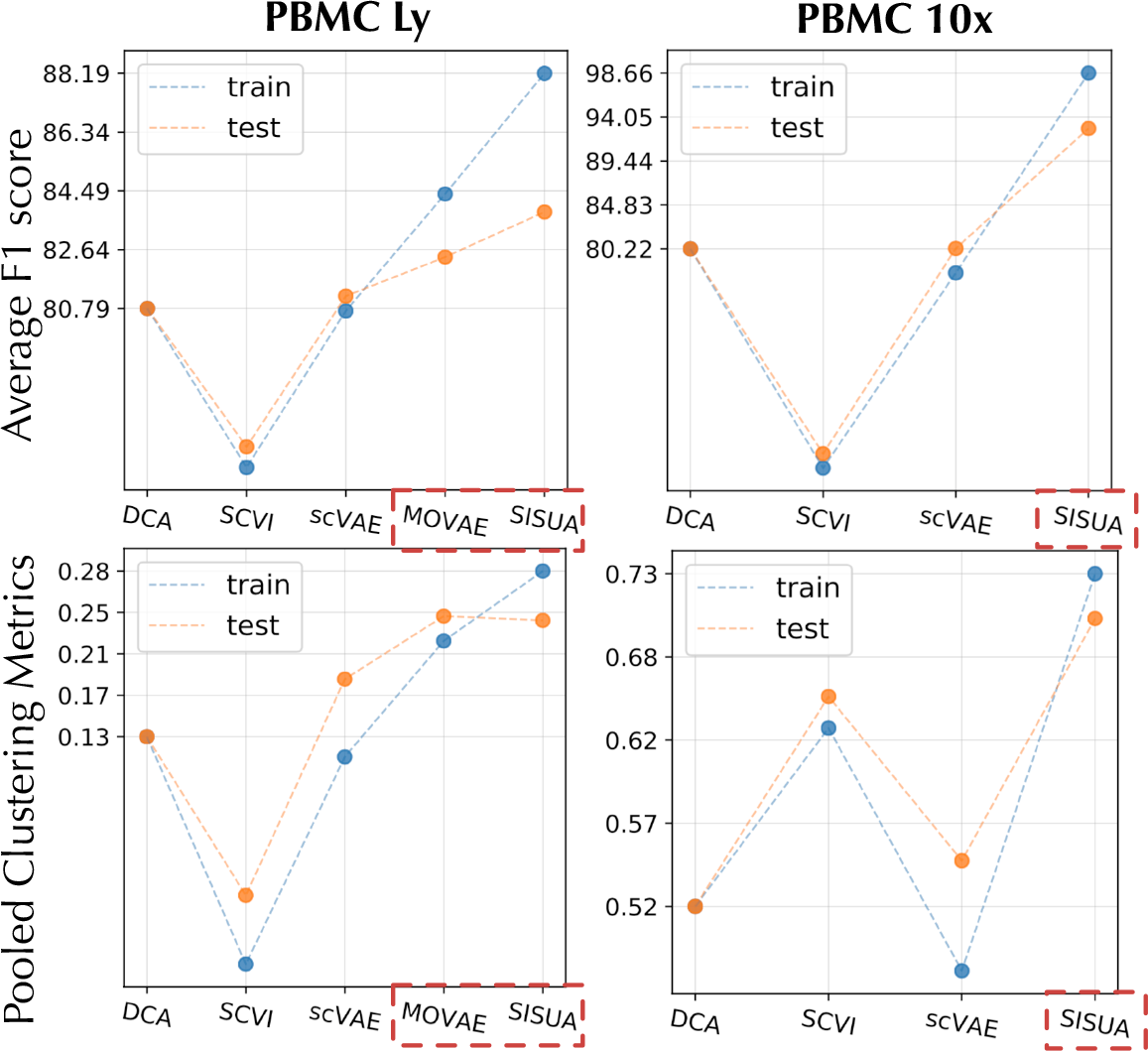
Latent spaces are evaluated by two benchmarks: the average of F1 scores from a streamline protein/cell types classifiers; the pooled clustering metric calculated by averaging 4 measurements: *ARI*, *ASW*, *NMI*, *UCA*. The results are reported for *PBMC Ly* and *PBMC 10x* datasets. The train results are shown in blue dots, the test results in orange dot. Semi-supervised models are highlighted by red dashed boxes.

**Table 1:** Marginal log likelihood for a held-out subset of each dataset, number of cells and genes also given for measuring the scale of each experiment.

| Dataset | PBMC Ly | PBMC 10x |
| --- | --- | --- |
| *scVAE* | -644.02 | -1494.75 |
| *MOVAE* | -643.83 | - |
| *SISUA* | **-642.70** | **-1494.51** |

### 5.3 Predictive protein distribution

Fig. 8 illustrates how the *SISUA* model has learned to predict protein marker levels. Dealing with continuous levels of markers requires powerful representation learning. The upper panel shows that the model has been able to learn the underlying structure that defines the surface protein levels cells to a high degree, visible as a high correlation of the predicted level and the ‘ground truth’ protein expression. However, there are many unexpected peaks that mismatching with the ‘ground truth’. The figure panels below attempt to explain this phenomenon. During the process of exploiting the correlation between gene expression and surface protein levels, the model learns to calibrate many faulty points in the given protein labels itself. When the latent space and the denoised space are colored by the ‘ground truth’ protein and the predicted protein level, the latter indicates more relevant structure, where many outliers for ‘CD4’ protein levels are cleaned and grouped into neat clusters (highlighted by green circles). It is notable that these are observed in both latent and denoised spaces, hence, *SISUA* has been able to capture relevant biological connections at multiple levels.

## 6 Generalizability

### 6.1 Goodness of fit on held-out data

The marginal log-likelihoods on the held-out data are presented in Table 1. We are able to compare fairly only *SISUA*, *scVAE* and *MOVAE*-models. In all cases, *SISUA* obtained the best fit. In the *PBMC 10x* dataset, with manually labeled cell types, the difference to completely unsupervised model was marginal but in the *PBMC Ly* subset, where independent protein data was used, a larger improvement was obtained.

**Figure 8:**
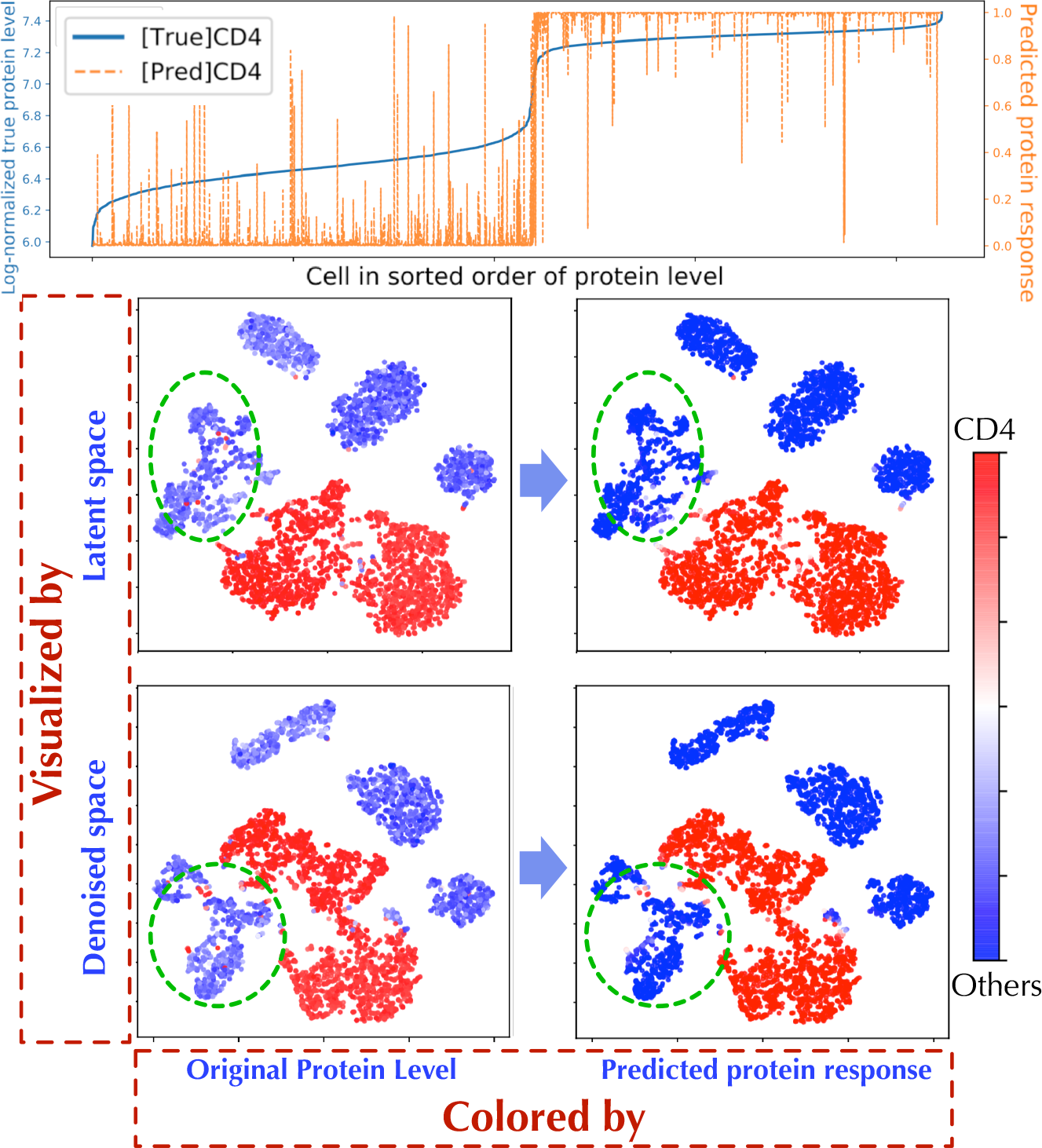
Comparing the original protein level and predicted protein response on *PBMC Ly* data set

### 6.2 Structural integrity of denoised space

To evaluate whether the denoised gene expression spaces still maintain the same essential variability model as the original data, we performed an imputation experiment shown in Fig. 9. The positions of the data points were projected into the total-variability space by PCA (trained on original data). To confirm that relevant biological information was preserved during the denoising processing, we inspected two properties: First, by coloring the cell types (first row) we could confirm whether similar cells still form clusters that reside in the same position in the total-variability space. Second, the cell size (colored in second row) was compared to that in the original data. We observed that *DCA* is altering both the cell types and cell size (blue circle), yet keeps a high amount of variability. *scVI* has very low variability, but performs good on cell type clustering and preserving cell sizes. *scVAE* provides better variability but slightly worse cell size. *SISUA* improves the variability compared to *scVAE*, and also improved the cell type model (red circles). Additionally, *SISUA* slightly increased the cell size compared to *scVAE* although this was not explicitly modeled.

## 7 Efficiency and scalability of semi-supervised training

### 7.1 Efficiency

In order to evaluate, whether the amount of labeled data available at the training phase affects the model quality, we measured the goodness in three different ways (Fig. 10); model fit in terms of the marginal log-likelihood, average F1 of protein/cell-type classifier in the latent space and average correlation between marker gene and the protein. We noticed in the PBMC Ly subset that adding only 1% of labels degraded the performance in all cases, while addition of 10% gave a clear boost in all three metrics. As expected, addition of labeled examples systematically improved the model in the PBMC Ly case. In the case of *PBMC 10x*, the situation is not as clear, since the average F1 improved until 80% of the training examples are labeled. However, the marginal log-likelihood does not show systematic behavior. One reason for this non-systematic behavior could be errors in *PBMC 10x* cell-type labels.

**Figure 9:**
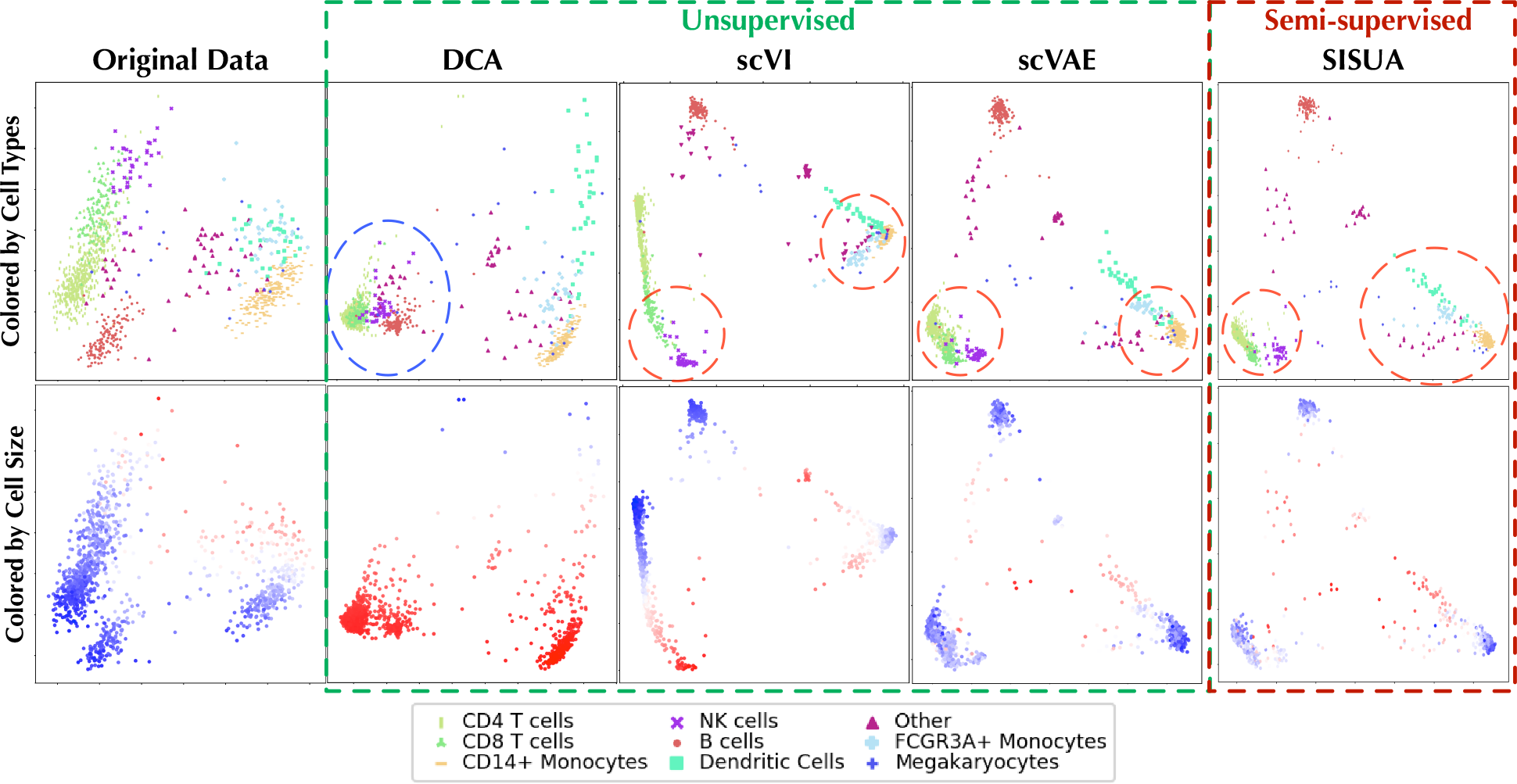
A PCA model trained using original gene expression data of the *PBMC 10x* data set was used to project the denoised gene expression from different models into its space. The top row is colored by cell type, the bottom row by denoised cell size (red color indicates large, white color mid-range and blue small cell size).

**Figure 10:**
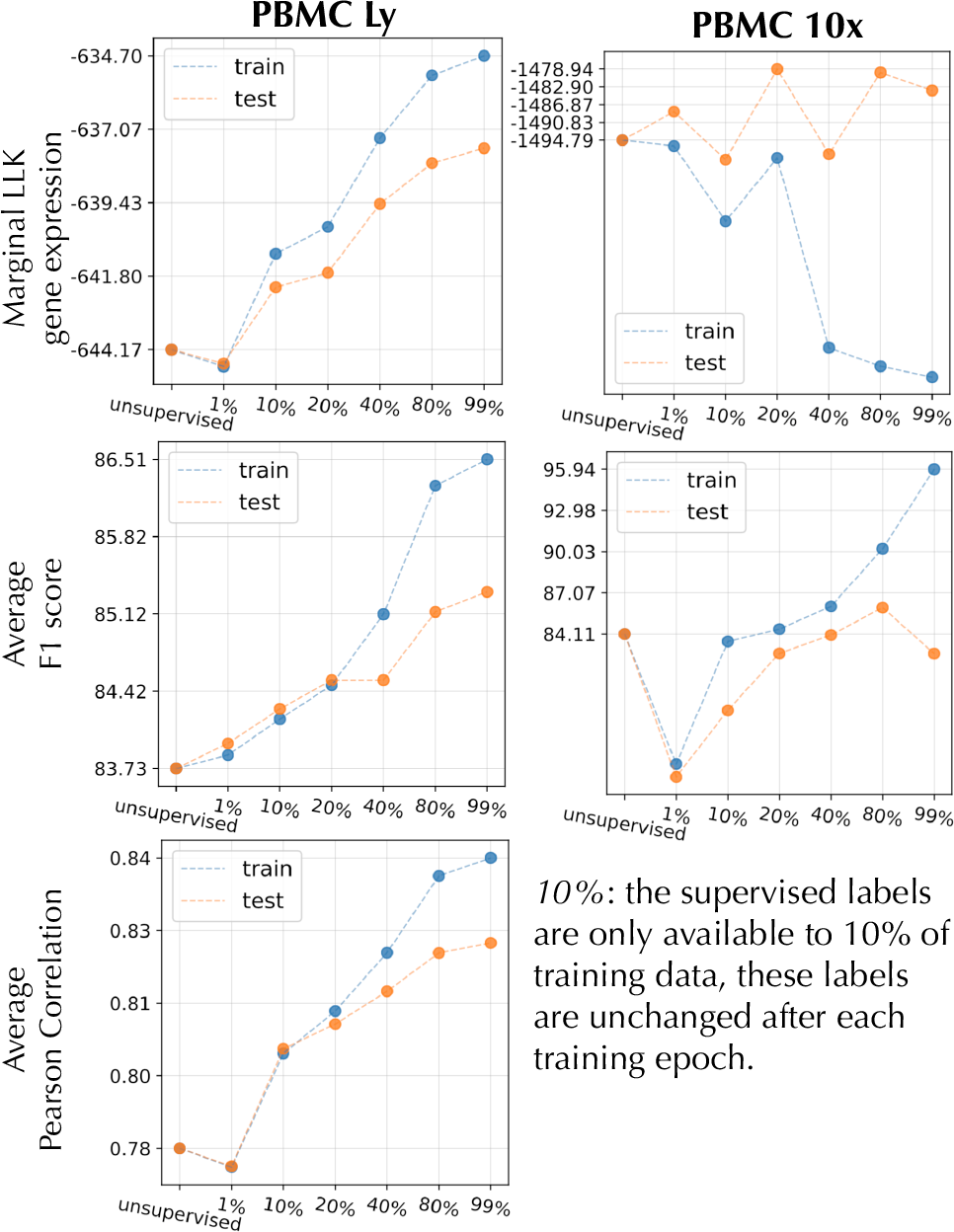
The performance on two data sets (*PBMC Ly* and *PBMC 10x*) is shown for different amount of labels utilized in training. The *X*-axis represents 7 systems with incresing amount of labeled data available for the semi-supervised objective (*Note:* no marker gene/protein pair is available for *PBMC 10x*).

**Figure 11:**
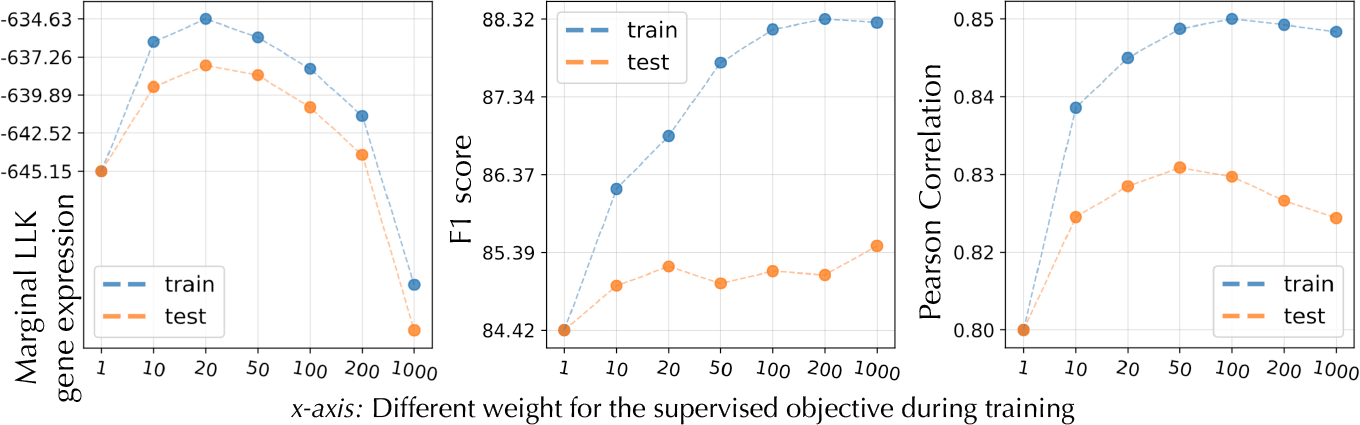
Performance of *SISUA* when γ (i.e. the weight of semi-supervised objective) is variated.

Fig. 11 emphasizes the important role of fine-tuning the γ parameter to balance the benefit of supervised learning and not over-powering the modeling of single-cell gene expression. We notice in all cases that γ of more than 20 gives clear improvement over the unsupervised case. However, when γ is increased more, the model starts to favour more the proteins than gene expression in the reconstruction.

### 7.2 Scalability

**Figure 12:**
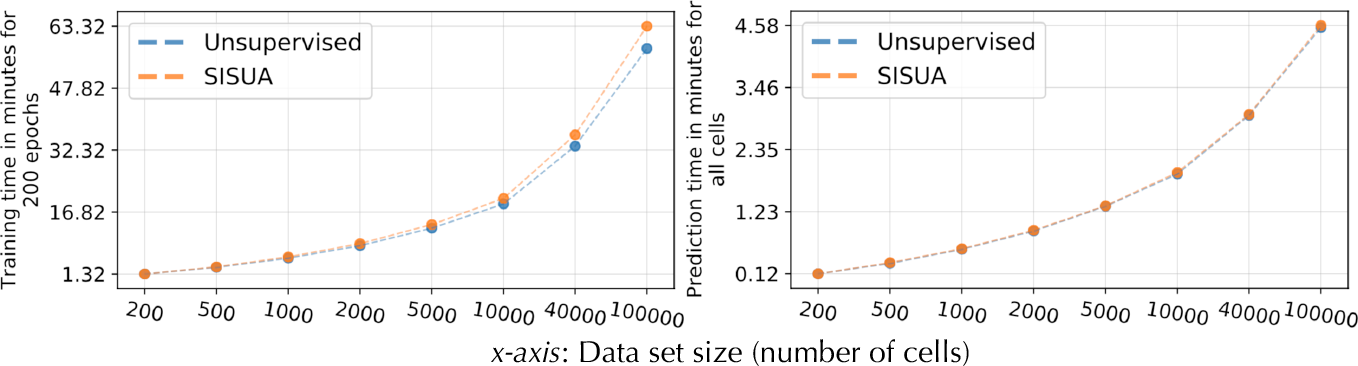
Running time for the training phase (left figure) and evaluation phase (right figure) for the unsupervised model and semi-supervised model (*SISUA*). The algorithms were ran on an eight-core Intel Xeon CPU E5-1630, and one NVIDIA GeForce GTX 1080.

Finally, the training algorithm running time as a function of number of cells is shown in the Fig. 12. We observed that the semi-supervised extension added a very minor increment to the running time when compared to the unsupervised variant. It is at most 8% longer compared to the unsupervised for 100000 cells.

It also is notable that the *SISUA* model introduces no extra running time during the evaluation phase, since no extra data is needed. Instead, we get the extra benefit of obtaining protein level predictions, better modeling of gene expression, and better representation of cell profiles in the latent space.

## 8 Conclusion

In classical machine learning, where the goal is to obtain accurate predictors, the semi-supervised approaches were motivated by the lack of labeled data. Unlabeled data is easy to obtain, but obtaining accurate labels is typically time consuming and expensive. In that respect, the unlabeled data is used to obtain higher accuracy predictive model. In the field of single-cell RNA-seq, we design a new model, *SISUA*, and turn this idea ‘upside down’, our task is to perform unsupervised analysis on the single-cell gene expression profiles and we are interested to see whether slight supervision can assist in producing biologically meaningful latent representations. Our results corroborate the merits of the semi-supervised extension. In addition to better interpreted latent representations, the method also enhances the imputation of RNA sequenced counts to more biologically meaningful places.

*SISUA* provides extra utility for labeling or predicting the cell types or surface protein levels of unseen data, this information has been proven to be valuable for cell diagnostic and analysis. Our research proposes general guidelines to implement an efficient and practical biological semi-supervised system. These policies are the substrate to develop more advanced designs and leverage the variety of external biological data that can benefit single-cell modeling.

## Acknowledgments

This research was partially funded by the CZI Collaborative Computational Tools for the Human Cell Atlas, Academy of Finland (grant #313970) and Finnish Scientific Advisory Board for Defence (MATINE) project nr. 2500M-0106. We gratefully acknowledge the support of NVIDIA Corporation with the donation of the Titan Xp GPU used for this research.

## Footnotes

1 Code and reproducibility: https://github.com/trungnt13/sisua

2 pbmc_10k_protein_v3_filtered_feature_bc_matrix.tar.gz

## References

[1] Fuchou Tang, Catalin Barbacioru, Yangzhou Wang, Ellen Nordman, Clarence Lee, Nanlan Xu, Xiaohui Wang, John Bodeau, Brian B Tuch, Asim Siddiqui, Kaiqin Lao, and M Azim Surani. mRNA-Seq whole-transcriptome analysis of a single cell. Nature Methods, 6:377–382, 2009.

[2] Byungjin Hwang, Ji Hyun Lee, and Duhee Bang. Single-cell RNA sequencing technologies and bioinformatics pipelines. Experimental & Molecular Medicine, 50(96), 2018.

[3] Eva Hedlund and Qiaolin Deng. Single-cell RNA sequencing: Technical advancements and biological applications. Molecular Aspects of Medicine, 59:36–46, 2018.

[4] Wei Vivian Li and Jingyi Jessica Li. An accurate and robust imputation method scimpute for single-cell RNA-seq data. Nature Communications, 9(997), 2018.

[5] David van Dijk, Roshan Sharma, Juozas Nainys, Kristina Yim, Pooja Kathail, Ambrose J. Carr, Cassandra Burdziak, Kevin R. Moon, Christine L. Chaffer, Diwakar Pattabiraman, Brian Bierie, Linas Mazutis, Guy Wolf, Smita Krishnaswamy, and Dana Pe’er. Recovering gene interactions from single-cell data using data diffusion. Cell, 174(3):716–729, 2018.

[6] Diane Lambert. Zero-inflated Poisson regression, with an application to defects in manufacturing. Technometrics, 34(1):1–14, 1992.

[7] Gary King. Variance specification in event count models: From restrictive assumptions to a generalized estimator. American Journal of Political Science, 33(3):762–784, 1989.

[8] Emma Pierson and Christopher Yau. Zifa: Dimensionality reduction for zero-inflated single-cell gene expression analysis. Genome biology, 16(1):1, 2015.

[9] D. Risso, F. Perraudeau, S. Gribkova, S. Dudoit, and J. Vert. A general and flexible method for signal extraction from single-cell rna-seq data. Nature Communications, 284(9), 2018.

[10] Romain Lopez, Jeffrey Regier, Michael B Cole, Michael Jordan, and Nir Yosef. Bayesian inference for a generative model of transcriptome profiles from single-cell rna sequencing. bioRxiv, 2018.

[11] Gökcen Eraslan, Lukas M. Simon, Maria Mircea, Nikola S. Mueller, and Fabian J. Theis. Single-cell rna-seq denoising using a deep count autoencoder. Nature Communications, 10(1):390, 2019.

[12] Yuval Hart, Hila Sheftel, Jean Hausser, Pablo Szekely, Noa Bossel Ben-Moshe, Yael Korem, Avichai Tendler, Avraham E Mayo, and Uri Alon. Inferring biological tasks using pareto analysis of high-dimensional data. Nature Methods, 12:233–235, 2015.

[13] Xiaojie Qiu, Arman Rahimzamani, Li Wang, Qi Mao, Timothy Durham, José L McFaline-Figueroa, Lauren Saunders, Cole Trapnell, and Sreeram Kannan. Towards inferring causal gene regulatory networks from single cell expression measurements. bioRxiv, 2018.

[14] Christopher Heje Grønbech, Maximillian Fornitz Vording, Pascal N Timshel, Casper Kaae Sønderby, Tune Hannes Pers, and Ole Winther. scvae: Variational auto-encoders for single-cell gene expression data. bioRxiv, 2018.

[15] Diederik P. Kingma, Danilo J. Rezende, Shakir Mohamed, and Max Welling. Semi-supervised learning with deep generative models. In Proceedings of the 27th International Conference on Neural Information Processing Systems, NIPS’14, pages 3581–3589, Cambridge, MA, USA, 2014. MIT Press.

[16] Marlon Stoeckius, Christoph Hafemeister, William Stephenson, Brian Houck-Loomis, Pratip K. Chattopadhyay, Harold Swerdlow, Rahul Satija, and Peter Smibert. Simultaneous epitope and transcriptome measurement in single cells. 14(9):865–868, Sep 2017.

[17] C.M. Bishop. Pattern Recognition and Machine Learning. Springer Science+Business Media, LLC, New York, 2006.

[18] M. E. Tipping and C. M. Bishop. Probabilistic principal component analysis. Journal of the Royal Statistical Society, Series B, 21:611–622, 1999.

[19] D. Rumelhart, G. Hinton, and R. Williams. Learning representations by back-propagating errors. Nature, 323:533–536, 1986.

[20] Andrew Gelman, John B. Carlin, Hal S. Stern, and Donald B. Rubin. Bayesian Data Analysis. Chapman and Hall/CRC, 2nd ed. edition, 2004.

[21] Rich Caruana. Multitask learning. Machine Learning, 1:41–75, July 1997.

[22] Trung Ngo Trong, Ville Hautamaki, and Kong Aik Lee. Deep language: a comprehensive deep learning approach to end-to-end language recognition. Odyssey: the Speaker and Language Recognition Workshop, 2016.

[23] Trung Ngo Trong, Kristiina Jokinen, and Ville Hautamäki. Enabling spoken dialogue systems for low-resourced languages: End-to-end dialect recognition for north sami. In Proceedings of the 9th International Workshop on Spoken Dialogue Systems Technology, Singapore, 2018.

[24] Antti Rasmus, Harri Valpola, Mikko Honkala, Mathias Berglund, and Tapani Raiko. Semi-supervised learning with ladder network. CoRR, abs/1507.02672, 2015.

[25] Ian Goodfellow, Jean Pouget-Abadie, Mehdi Mirza, Bing Xu, David Warde-Farley, Sherjil Ozair, Aaron Courville, and Yoshua Bengio. Generative adversarial nets. In Z. Ghahramani, M. Welling, C. Cortes, N. D. Lawrence, and K. Q. Weinberger, editors, Advances in Neural Information Processing Systems 27, pages 2672–2680. Curran Associates, Inc., 2014.

[26] Diederik P. Kingma and Jimmy Ba. Adam: A method for stochastic optimization. CoRR, abs/1412.6980, 2014.

[27] Douglas Reynolds. Universal Background Models, pages 1349–1352. Springer US, Boston, MA, 2009.

[28] Paulina Hensman and David Masko. The impact of imbalanced training data for convolutional neural networks. Degree Project in Computer Science, KTH Royal Institute of Technology, 2015.

[29] Alexandre Dalyac, Prof Murray Shanahan, and Jack Kelly. Tackling class imbalance with deep convolutional neural networks. Thesis, Imperial College London, 2014.

[30] Grace X.Y. Zheng, Jessica M. Terry, and et al. Massively parallel digital transcriptional profiling of single cells. bioRxiv, 2016.

